# Nuclear actin is required for transcription during *Drosophila* oogenesis

**DOI:** 10.1101/358028

**Authors:** Maria Sokolova, Henna M. Moore, Bina Prajapati, Joseph Dopie, Leena Meriläinen, Mikko Honkanen, Rita Cerejeira Matos, Minna Poukkula, Ville Hietakangas, Maria K. Vartiainen

## Abstract

Actin influences gene expression at multiple levels. It regulates the activity of specific transcription factors, such as myocardin-related transcription factor (MRTF), is a component of many chromatin remodelers and linked to transcription by all three RNA polymerases (Pol). However, the molecular mechanisms by which actin participates in the gene-specific vs. general transcription have remained unclear. Here we use chromatin immunoprecipitation followed by deep sequencing (ChIP-seq) in *Drosophila* ovaries to demonstrate that binding of actin to the *Act5C* gene is not dependent on the Mrtf transcription cofactor. At the genome-wide level, actin interacts with essentially all transcribed genes and co-occupies most gene promoters together with Pol II. On highly expressed genes, actin and Pol II can be found also on the gene bodies. Manipulation of nuclear transport factors for actin leads to decreased expression of egg shell genes, demonstrating the *in vivo* relevance of balanced nucleo-cytoplasmic shuttling of actin for transcription.

## Introduction

In addition to its essential roles as part of the cytoskeleton, actin also regulates gene expression in the nucleus. Actin is a component of many chromatin remodeling complexes [reviewed by (Kapoor and Shen, 2013)] and linked to transcription by all three RNA polymerases (Hofmann et al., 2004; Hu et al., 2004; Philimonenko et al., 2004). Actin seems to have a positive role on general transcription, since reduced availability of nuclear actin, due to either inhibition of the active nuclear import of actin (Dopie et al., 2012), activation of a mechanosensory complex consisting of emerin, non-muscle myosin II and actin (Le et al., 2016) or polymerizing nuclear actin into stable filaments (Serebryannyy et al., 2016), attenuates transcription. Nevertheless, the exact mechanism and the *in vivo* relevance of this process have remained unclear. Actin also negatively regulates the transcription of specific genes. For example, actin regulates both the nuclear localization and activity of myocardin related transcription factor A (MRTF-A; also known as MAL/MKL1), which is cofactor of the essential transcription factor SRF (Miralles et al., 2003; Vartiainen et al., 2007). Actin monomer-binding prevents MRTF-A from activating SRF in the nucleus. This regulation has been postulated to take place at the level of target genes (Vartiainen et al., 2007), but how the opposing effects of actin on transcription are resolved on chromatin is not obvious. Moreover, the genome-wide binding pattern of actin in the context of RNA polymerase II (Pol II) mediated transcription has remained elusive. Importantly, actin itself is one of the target genes for SRF (Salvany et al., 2014), generating a feedback loop, where actin levels are controlled by the actin dynamics cycle. Here we show that chromatin-binding of actin is not dependent on Mrtf transcription factors and that, at the genome-wide level, actin interacts with essentially all transcribed genes in *Drosophila* ovaries, with a pattern depending on the expression level of the gene. Finally, we demonstrate the functional relevance of nuclear actin for gene transcription *in vivo*.

## Results and discussion

### Actin is involved in transcription of *Act5C* independently of Mrtf

To clarify the role of actin in general vs. gene-specific transcriptional regulation, we examined actin-chromatin interactions in *Drosophila* ovaries, where Mrtf has been shown to regulate *Act5C* transcription (Salvany et al., 2014). We performed chromatin immunoprecipitation followed by deep sequencing (ChIP-seq) of Mrtf-GFP, actin and Pol II phosphorylated at serine 5 (Pol II S5P) in ovaries of wild type (*w*^1118^) and *Mrtf* mutant (*mal-d^Δ7^*) flies, where *Mrtf* expression is abolished (Somogyi and Rorth, 2004), as well as in flies ubiquitously expressing GFP-tagged version of Mrtf (*tub mal-d3xGFP*) (Salvany et al., 2014) (**Figure 1A**). Deletion and overexpression of Mrtf displayed decreased and increased expression of *Act5c*, respectively (**Figure 1B**), and Mrtf bound to promoter and upstream region of the *Act5C* gene (**Figure 1A,C,D**), in agreement with previous studies (Salvany et al., 2014). Pol II S5P bound to transcription start sites of *Act5C* in all three fly strains (**Figure 1A,C,D**). Interestingly, the binding pattern of actin was different than that of Mrtf, and a substantial actin signal was found on the gene body of the *Act5C* gene (**Figure 1A,E**). Importantly, actin signal was not reduced in *mal-d^Δ7^* flies (**Figure 1F**), indicating that actin-binding to the *Act5C* gene is not dependent on Mrtf.

**Figure 1.**
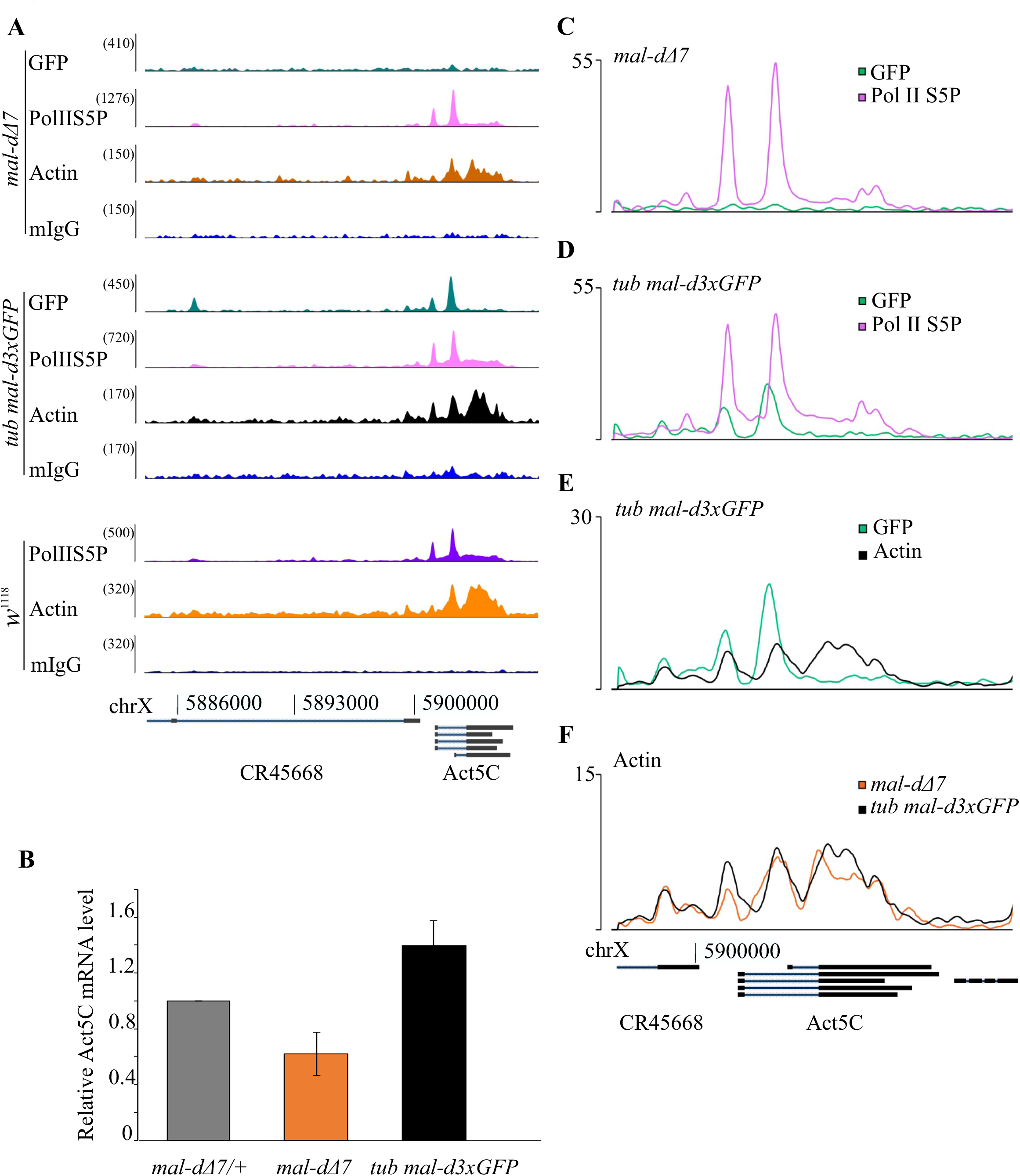
Actin-binding to the *Act5C* gene is not dependent on Mrtf. A. ChIP-seq analysis of Mrtf-GFP, actin and Pol II S5P at the *Act5C* gene region on chromosome X. Fly strains and antibodies used are indicated on the left, and signal intensity as number of reads is shown above each track; actin and the control antibody IgG are shown on the same scale. B. mRNA levels of Act5C in the indicated fly strain measured by qPCR. Rpl32 was used as internal control, data is normalized to mal-dΔ7/+ and is mean from two independent measurements with standard deviation. C,D. Binding profile of Pol II S5P (purple) and Mrtf-GFP (green) on *Act5C* gene in ovaries from *mal-dΔ7* (C) and *tub mal-d3xGFP* (D) fly strains. Read counts are normalized to inputs. E. Binding profile of actin (black) and Mrtf-GFP (green) on the Act5C gene in ovaries from *tub mal-d3xGFP* fly strain. Read counts are normalized to inputs. F. Binding profile of actin on the *Act5C* gene in ovaries from *tub mal-d3xGFP* (black) and in *mal-dΔ7* (light brown) flies.

### Actin interacts with transcribed genes with a pattern depending on their expression level

Further ChIP-Seq analysis of the *w*^1118^ fly strain revealed actin on the promoters of essentially all transcribed genes together with Pol II S5P (**Figure 2A**). Peak-calling confirmed the substantial overlap between actin and Pol II S5P binding sites (**Figure 2B**). However, detailed analysis showed that actin binds promoters slightly before the transcription start site (TSS) and Pol II S5P enrichment (**Figure 2C**), indicating that actin could be involved in transcriptional initiation, perhaps via pre-initiation complex formation, as suggested before (Hofmann et al., 2004).

**Figure 2.**
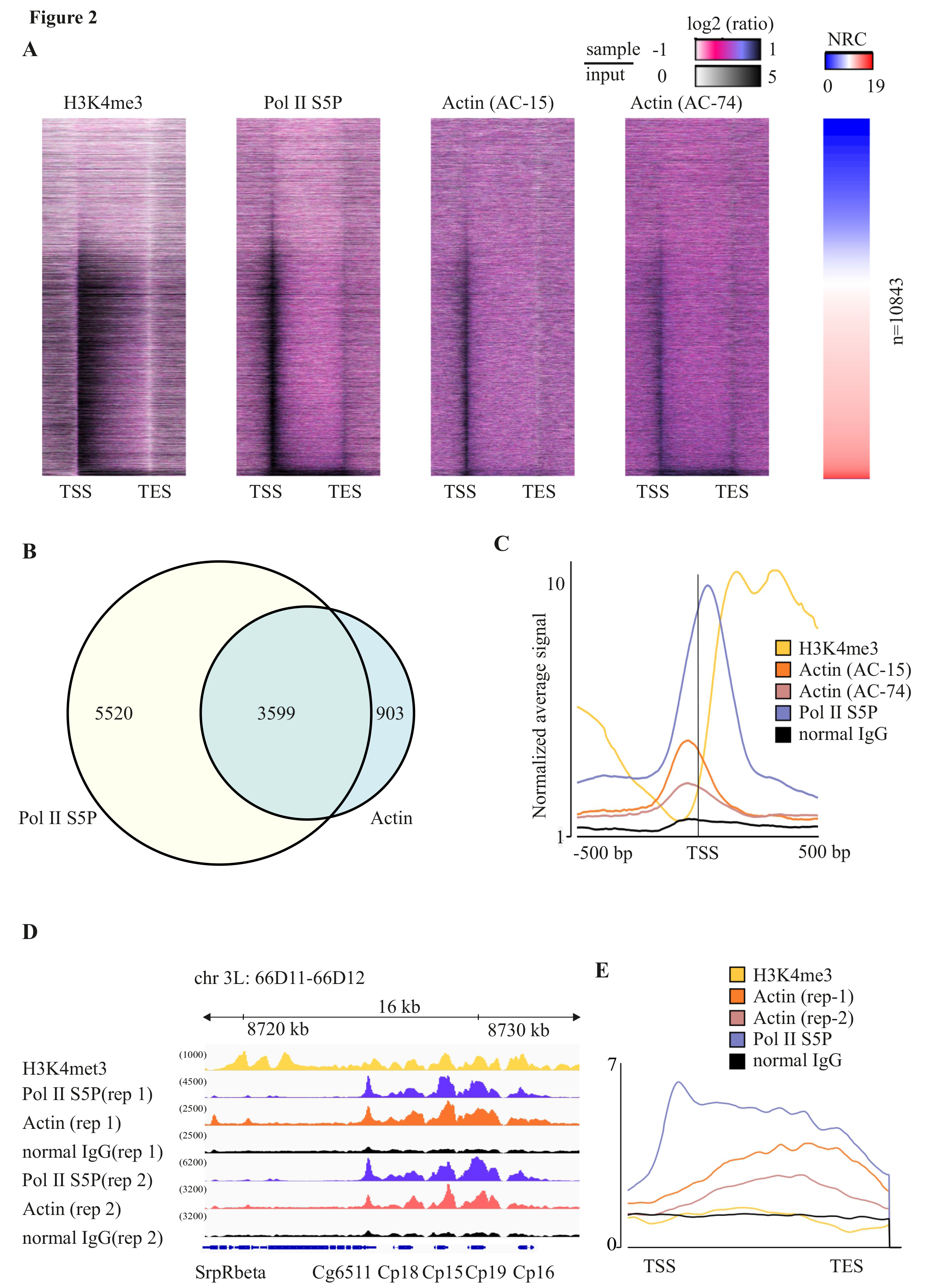
Actin colocalizes with Pol II at TSS and gene bodies of transcribed genes. A. Heatmap of the ratio between the sample (histone H3K4met3, Pol II S5P, and actin with two antibodies, AC-74 and AC-15) and input ChIP-Seq signals across gene regions, standardized and segmented into 200 bins. Transcription start sites (TSS) and transcription end sites (TES) are indicated. Genes are sorted according to normalized read count (NRC) of RNA-seq data from *w*^1118^fly ovaries (right panel). B. Venn diagram showing overlap of actin (AC-74) and Pol II S5P peaks from ChIP-seq. C. Average signal of read counts normalized to the input from −500 bp to +500 bp from the TSS of gene loci (n=10843). D. Binding profile of actin and Pol II on *chorion* genes at 66D locus of chromosome 3L. Antibodies used in ChIP-Seq are indicated on the left, and signal intensity as number of reads is shown in parentheses above each track. Results from two experiment replicates (rep) are shown. E. ChIP-seq with the indicated antibodies with average signal of read counts normalized to input shown across the gene body of known eggshell protein encoding genes (Tootle et al., 2011).

Similarly to the *Act5C* gene (**Figure 1**), actin was also found, together with Pol II S5P, on gene bodies of certain genes (**Figure 2A**, genes at the bottom have highest expression). These included, for example, the highly transcribed *chorion* genes (**Figure 2D**) involved in eggshell formation. On these genes actin is enriched more towards the transcription end site (TES) than the TSS (**Figure 2E**). Notably, both actin antibodies produced a very similar binding pattern on chromatin (**Figure 2A,B,D,E**). This genome-wide analysis shows that actin interacts with most transcribed genes in *Drosophila* ovaries, and that depending on the expression level of the gene, actin can be found both on the promoters and gene bodies. This data can thus consolidate previous ChIP studies of actin that have reported variable binding to different genomic sites depending on the specific gene analyzed (Hu et al., 2004; Obrdlik et al., 2008; Philimonenko et al., 2004; Ye et al., 2008). Whether the binding pattern of actin reflects its dual roles in transcription, both during transcription initiation and elongation, or whether the recruitment to gene bodies represents a specific requirement for actin upon high transcriptional activity, awaits further studies. An obvious candidate for recruiting actin to the genes is Pol II, which based on our ChIP-seq studies co-occupies most actin-binding sites (**Figure 2**), although not with exactly the same pattern. Other candidates include the different chromatin remodeling complexes containing actin (Kapoor and Shen, 2013), as well as the elongation factor P-TEFb (Qi et al., 2011).

### Active transport of nuclear actin is required for egg shell gene transcription

To study if active maintenance of nuclear actin levels is required for transcription in *Drosophila* ovaries similarly as in mammalian cells (Dopie et al., 2012), we generated a mutant of the nuclear actin import receptor, RanBP9 (*Drosophila* orthologue of Importin-9) (**Figure 3A**; see also Materials and methods). Similarly to Importin-9 knockdown in mammalian cells (Dopie et al., 2012), loss of RanBP9 in *Drosophila* resulted in decreased nuclear actin levels (**Figure 3B,C**), while the total actin levels were not significantly altered (**Figure 3D**). On the same genetic background, the *RanBP9*^Δ1^ mutants were viable, but females laid fewer eggs than control flies (**Figure 3E**), and these eggs failed to develop.

**Fig. 3.**
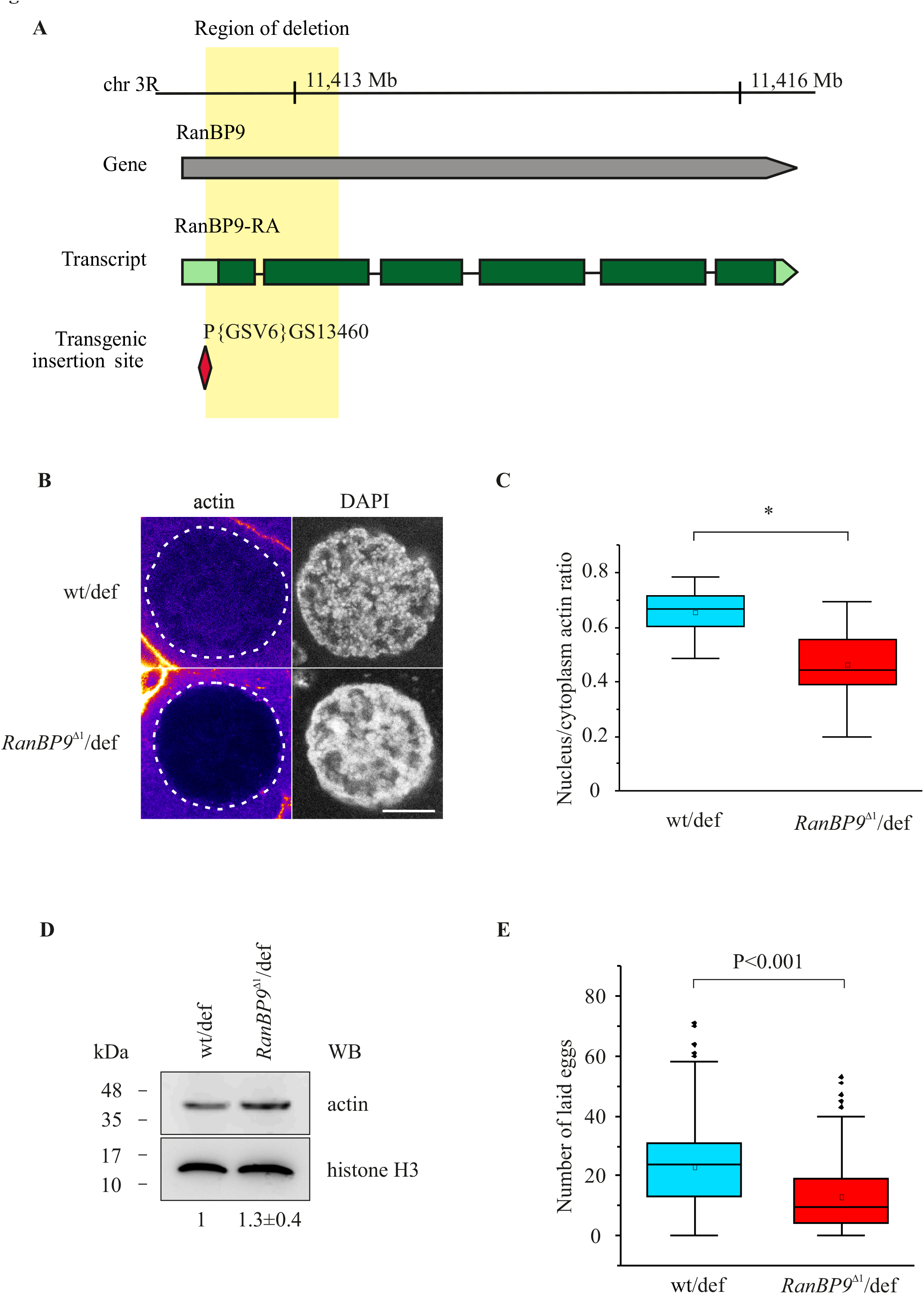
Generation of RanBP9 mutant fly with decreased nuclear actin. A. Schematic of the RanBP9 locus. The region of deletion (light yellow) generated by imprecise excision of P{GSV6}GS13460. B. Confocal microscopy images of nurse cell nuclei of ovarian egg chambers stained with actin antibodies and DAPI. Scale bar 10 µm. C. Quantitation of nucleus to cytoplasm ratio of actin staining intensities in nurse cells. Data is from three independent experiments with N=32 (wt/def) and N=29 (*RanBP9*^Δ1^/def). Mann-Whitney Test, p<0.05. Boxes represent 25-75% and the error bars range within 1.5IQR. The line in the middle is median and the open square is mean. D. Western blots from the whole fly lysates probed with anti-actin antibody. Quantitation of actin amount (below the blots) is from three independent experiments with wt/def normalized to 1 and ± representing SD. No significance by Student’s T test. E. Numbers of eggs laid by the indicated flies. N=289 (wt/def) and N=214 (*RanBP9*^Δ1^/def) from six independent experiments. Student’s t-test, p<0.001. Data shown as in C. Black diamonds are outliers.

In contrast to our previous results from mammalian cells, RNA-seq analysis of the *RanBP9*^Δ1^ mutant ovaries did not reveal dramatic transcriptional downregulation upon inhibiting active nuclear import of actin (**Figure 4A and Supplementary table 1**). We note that in mammalian cells, Importin-9 depletion led to a greater reduction in nuclear actin levels (Dopie et al., 2012) than the *RanBP9*^Δ^ deletion reported here (**Figure 3C**). Whether the fly utilizes additional nuclear import mechanisms for actin or whether the underlying biological complexity creates differential sensitivity to nuclear actin levels remains to be determined. Nevertheless, several genes encoding for chorion proteins showed reduced expression in the *RanBP9*^Δ1^ compared to control (marked as red in **Figure 4A)**, and RT-qPCR confirmed the significant downregulation for a subset of them (**Figure 4B)**. Importantly, the same transcripts showed reduced expression also when RanBP9 expression was silenced by RNAi specifically in the follicle cells (**Figure 4C**), which are the cells that express the *chorion* genes to deposit the eggshell over the oocyte. Since RanBP9 could also have other import cargoes than actin, we used overexpression of Exportin 6, the nuclear export receptor for actin (Stuven et al., 2003), as an alternative method to manipulate nuclear actin in follicle cells. Also this led to reduction in *chorion* gene expression (**Figure 4C**), further supporting the notion that balanced nuclear transport of actin is required for appropriate transcription of egg shell genes. Finally, the eggs laid by the *RanBP9*^Δ1^ females displayed morphologically abnormal (**Figure 4D**) and short (**Figure 4E**) dorsal appendages, which are specialized structures of the egg shell used by the embryo for breathing. Deregulated *chorion* gene expression thus has phenotypic consequences and could explain why the eggs laid by the *RanBP9*^Δ1^ females do not develop.

**Fig. 4.**
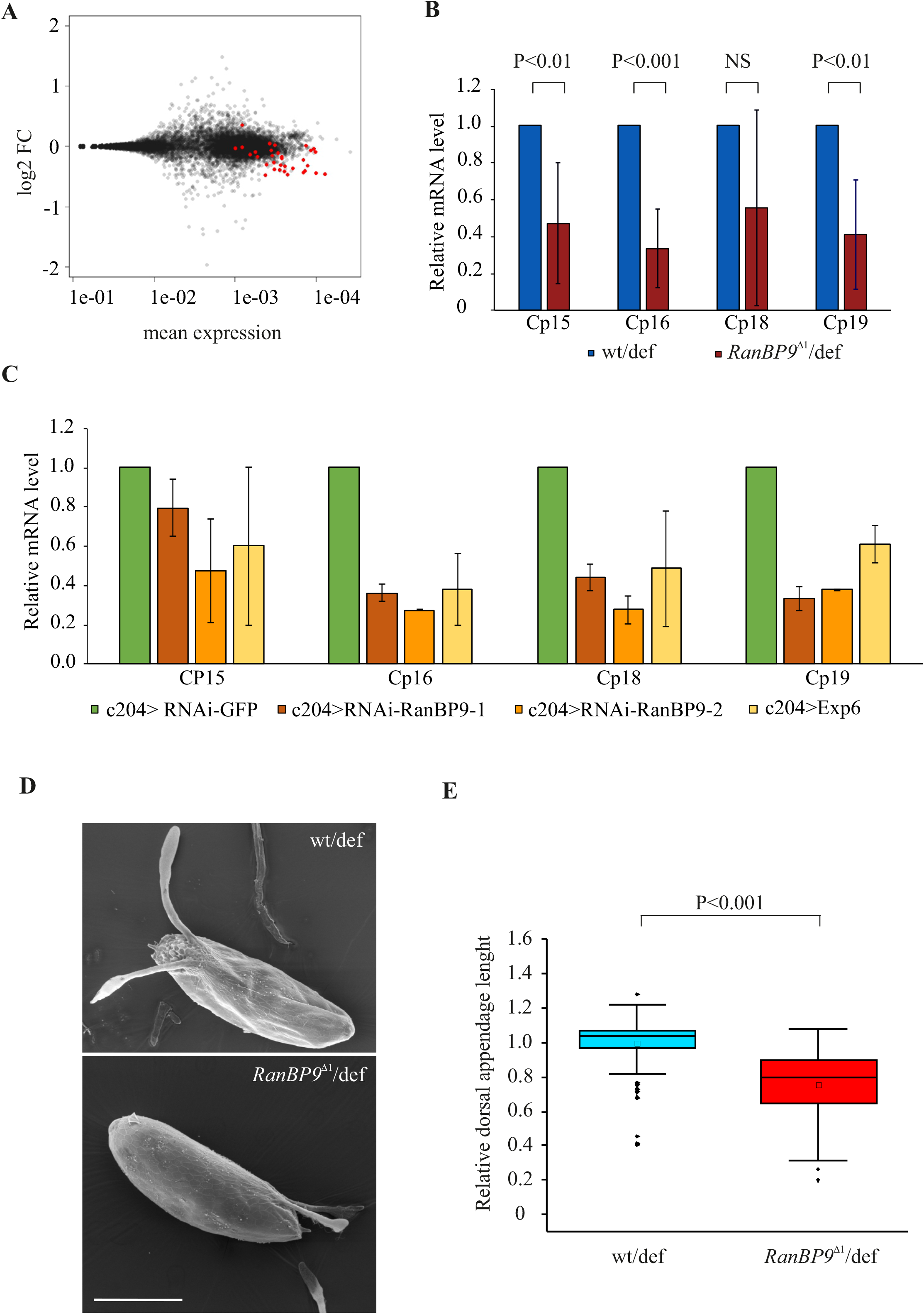
RanBP9 mutants display decreased expression of chorion protein genes and defective egg shell formation. A. MA-plot of RNA-Seq data. The transcripts of known eggshell proteins are indicated in red. B. Relative expression of 4 chorion protein transcripts in wt/def and *RanBP9*^Δ1^/def fly ovaries from five independent experiments. Data is normalized to wt/def. Statistics with student’s t-test. Error bars represent +/- SD. C. Relative expression of 4 chorion protein transcripts in the indicated fly strains from two independent experiments. Data is normalized to c204>RNAi-GFP and error bars represent +/- SD. D. Scanning electron micrographs of fly eggs with dorsal appendages. Representative images of control (wt/def) and *RanBP9*^Δ1^/def eggs are shown. Magnification 450x. Scale bar 200 µm. E. Relative lengths of dorsal appendages from eggs of indicated fly strains. Data is normalized to wt/def. N=91 (wt/def) and N=120 (*RanBP9*^Δ1^/def) from three independent experiments. Student’s t-test, p<0.001. Data shown as in 3C.

Taken together, these results enforce the importance of actin for transcription by showing that it interacts with virtually all genes transcribed by Pol II and that its balanced nuclear transport is required for transcription *in vivo*. Further studies are required to elucidate the molecular machineries that recruit actin both to the promoters and gene bodies.

## Materials and methods

### Antibodies

Primary antibodies used for the ChIP-seq included actin [AC-74 (A2228) and AC-15 (A1978), Sigma-Aldrich], Anti-RNA polymerase II CTD repeat YSPTSPS (phospho S5) (4H8; ab5408, Abcam), Anti-RNA polymerase II CTD repeat YSPTSPS (phospho S2) (ab5095, Abcam), Histone H3 (tri methyl K4) (ab8580, Abcam) and normal mouse IgG (sc-2025, Santa Cruz Biotechnology); for immunofluorescence anti-actin (A2103, Sigma-Aldrich); for WB anti-histone H3 (H0164, Sigma-Aldrich) and anti-actin [AC-15 (A1978), Sigma-Aldrich].

### Fly strains

All flies were maintained at +25°C. Fly strains from Bloomington Drosophila Stock Center included *w* 1118 (#3605), tub mal3xGFP (#58443), mal-dΔ7 (#58418), Df(3R)BSC469 (#24973), c204 (#3751) VALIUM20-EGFP.shRNA.1 (# 41556), HMS00804 (RNAi-RanBP9-1, # 33004), HMS00805 (RNAi-RanBP9-2 # 33005). P{GSV6}GS13460 (#205564) was from Kyoto Stock Center. UASp-Exp6 was a kind gift from Joachim Urban.

The *RanBP9*^Δ1^ mutant was generated from P{GSV6}GS13460 line using the method of imprecise excision. The mutants were screened with the following primers:

RanBP9_FW: 5’ TCGATTACTATCCAATCGTAA

RanBP9_RV: 5’ CACATGCGCACCGTGAGCTCC

The deletion was sequenced with the same primers and consisted of 839 bp deletion from 5’UTR to the end of second exon of RanBP9. This was further confirmed by RNA-seq. To minimize the influence of genetic background, we then crossed *w*^1118^ and *RanBP9*^Δ1^ flies with Df(3R)BSC469 deficiency stock (deletion of 86D8-87A2, which contains the *RanBP9* gene), and the resulting *RanBP9*^Δ1^/Df(3R)BSC469 (*RanBP9*^Δ1^/def) and control *w*^1118^/Df(3R)BSC469 (wt/def) fly lines were used in all experiments. The ability of the female flies to lay eggs was assessed by placing virgin females of each genotype with the same number of *w* 1118 males. On the day 4, flies were transferred to fresh vials with one male and one female in each vial. The total number of eggs produced over 24 h by each female was counted for 289 wt/def and 214 *RanBP9*^Δ1^/def females from six independent experiments.

The length of dorsal appendages were recorded either from live or frozen embryos that were laid by 5 day old wt/def and *RanBP9*^Δ1^/def females mated with *w*^1118^ males by using FLoid imaging station (Life Technologies) with Plan Fluorite 20x/0.45 objective and quantified with Fiji/ImageJ (Schindelin et al., 2012).

### ChIP-seq

For chromatin immunoprecipitation (ChIP) ovaries dissected from fly strains w1118, tub mal3xGFP and mal-dΔ7 were fixed in 1% paraformaldehyde/PBS for 10 min at RT, crosslinking was stopped by adding glycine to a final concentration of 0.125 M for five min, followed by homogenization using a pestle in 300 μl of RIPA buffer, and sonication with Bioruptor (Diagenode; number of cycles = 15, power = HIGH, ON = 30 sec, OFF = 30 sec). At least 100 ovaries were used per one IP. IPs were carried out with 5 µg antibody overnight at 4°C in a rotating wheel. The immuno-complexes were collected with 50 μl of protein A sepharose (17-0780-01, GE Healthcare) at 4 °C for two hours with rotation. For MAL-GFP immunoprecipitation, 50 µL of GFP beads (GFP-Trap ChromoTek) was used. The beads were pelleted by centrifugation at 4 °C for one minute at 500g and washed sequentially for five minutes on rotation with 1 ml of the following buffers: low-salt wash buffer (RIPA) (10 mM Tris–HCl (pH 8.0), 0.1% SDS, 1% Triton X-100, 1 mM EDTA, 140 mM NaCl, 0,1% sodium deoxycholate), high-salt wash buffer (10 mM Tris–HCl (pH 8.1), 0.1% SDS, 1% Triton X-100, 1 mM EDTA, 500 mM NaCl, 0,1% sodium deoxycholate) and LiCl wash buffer (10 mM Tris–HCl (pH 8.1), 0.25 mM LiCl, 0,5% IGEPAL CA-630, 0,5% sodium deoxycholate, 1 mM EDTA). Finally, the beads were washed twice with 1 ml of TE buffer (10 mM Tris–HCl (pH 8.0), 1 mM EDTA). Chromatin was eluted in 150 µl of 1% SDS in TE buffer. The cross-linking was reversed by adding NaCl to final concentration of 200 mM and incubating at 65 °C overnight. The eluate was treated with Proteinase K and the DNA was recovered by extraction with phenol/chloroform/isoamyl alcohol (25/24/1, by vol.) and precipitated with 0.1 volume of 3 M sodium acetate (pH 5.2) and two volumes of ethanol using glycogen as a carrier.

ChIP libraries were prepared for Illumina NextSeq 500 using NEBNext ChIP-Seq DNA Sample Prep Master Mix Set for Illumina (NEB E6240) and NEBNext^®^ Multiplex Oligos for Illumina^®^ (Index Primers Set 1) (NEB E7335) according to the manufacturer’s protocols. Sequencing was performed with NextSeq500 at Biomedicum Functional Genomics Unit (FuGU). ChIP-seq was performed in triplicate.

ChIP-Seq data sets were aligned using Bowtie2 [using Chipster software (Kallio et al., 2011)] to version dm6 (BDGP6.87) of the fly genome with the default settings. Peak calling was performed in R using BayesPeak package (Cairns et al., 2011; Spyrou et al., 2009). To visualize and present ChIP-seq data, we used Integrative Genomics Viewer [IGV; (Robinson et al., 2011)] and EaSeq (http://easeq.net) (Lerdrup et al., 2016).

### Immunofluorescence, microscopy and Western blotting

For immunostaining at least 10 pairs of ovaries were dissected in PBS containing protease inhibitors (complete Tablets EDTA-free, EasyPack, Roche), and fixed with 4% formaldehyde (EM grade, Electron Microscopy Sciences, 15710) for 20 minutes, at RT. Tissues were permeabilized in 0.5% Triton X-100 in PBS (PBT) for 10 minutes, and blocked with 4% bovine serum albumin (BSA) in PBT for 1 h. The ovaries were incubated with anti-actin antibodies (A2103, Sigma-Aldrich) for 2 nights at +4 °C. Tissues were washed in PBT containing 1% of BSA and incubated with the corresponded goat anti-rabbit Alexa Fluor™ 488 and DAPI for 3 hours at RT. Tissues were mounted in Prolong Gold (Molecular Probes, Life Technology). Images of egg chambers were acquired with Leica TCS SP8 confocal microscope equipped with an HC PL APO 93x/1.30 objective. Diode 405 and Argon laser lines were used for excitation. Image acquisition was performed with LASX software. For optimal nuclear imaging the pinhole was set as 1, and line averaging to 8. Fluorescent intensities of nuclei and cytoplasm were measured using Fiji/ImageJ software (Schindelin et al., 2012). The ratios of nucleus to cytoplasm intensities were calculated for each nurse cell from three independent experiments.

For Western blotting, lysates were prepared from wt/def and *RanBP9*^Δ1^/def females, five flies per sample in each experiment. Samples were prepared in Laemmli sample buffer and processed by SDS-12% PAGE for immunoblot analysis using antibodies: anti-histone H3 (H0164, Sigma-Aldrich) and anti-actin (AC-15 (A1978), Sigma-Aldrich).

### RNA-seq

Total RNA was extracted from dissected ovaries in triplicates from 6 wt/def and *RanBP9*^Δ1^ /def females with TRIzol (15596026, Thermofisher) according to the manufacturer’s protocol. Libraries were prepared Illumina NextSeq 500 using Ribo-Zero rRNA Removal Kit (Illumina) and the NEBNext Ultra Directional RNA Library Prep at the Biomedicum Functional Genomics Unit (FuGU) according to the manufacturer’s protocols.

RNA-seq data sets were aligned using TopHat2 (Kim et al., 2013) [using Chipster software (Kallio et al., 2011)] to version dm6 (BDGP6.87) of the fly genome with the default settings. Counting aligned reads per genes were performed with HTSeq (Anders et al., 2015). Differential expression analysis was performed with DESeq (Love et al., 2014). List of the transcribed genes (n=10843) in ovaries was based on the aligned reads count cutoff >1 from the RNA-Seq data from wt/def ovaries (Supplementary table X).

### Quantitative PCR (qPCR)

The wt/def and *RanBP9*^Δ1^/def females crossed with *w*^1118^ males were maintained at 25°C. The follicle cells specific GAL4 (c204) and UASp (EGFP-RNAi, RNAi-RanBP9-1, RNAi-RanBP9-2 and Exp6) fly strains were maintained at 25°C and transferred to 28°C after the cross.

Ovary samples from 5 day old females were used for quantitative comparison (5 pairs of ovaries per genotype in each experiment). RNA was prepared using NucleoSpin RNA Macherey-Nagel kit including DNase treatment (740955), and 0,5 μg of total RNA was used for reverse transcription and first strand cDNA synthesis with Maxima First Strand cDNA Synthesis Kit for RT-qPCR(K1641, Thermo Scientific). Quantitative PCR was performed using Bio-Rad Real-Time PCR Detection Systems CFX96. Chorion protein transcripts levels were calculated relative to reference gene Rpl32. Primers used are listed below:

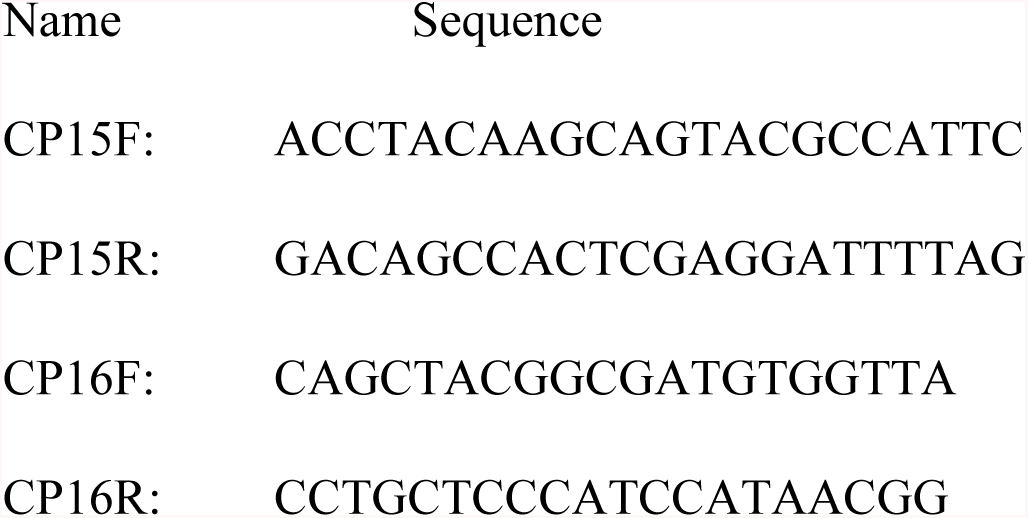
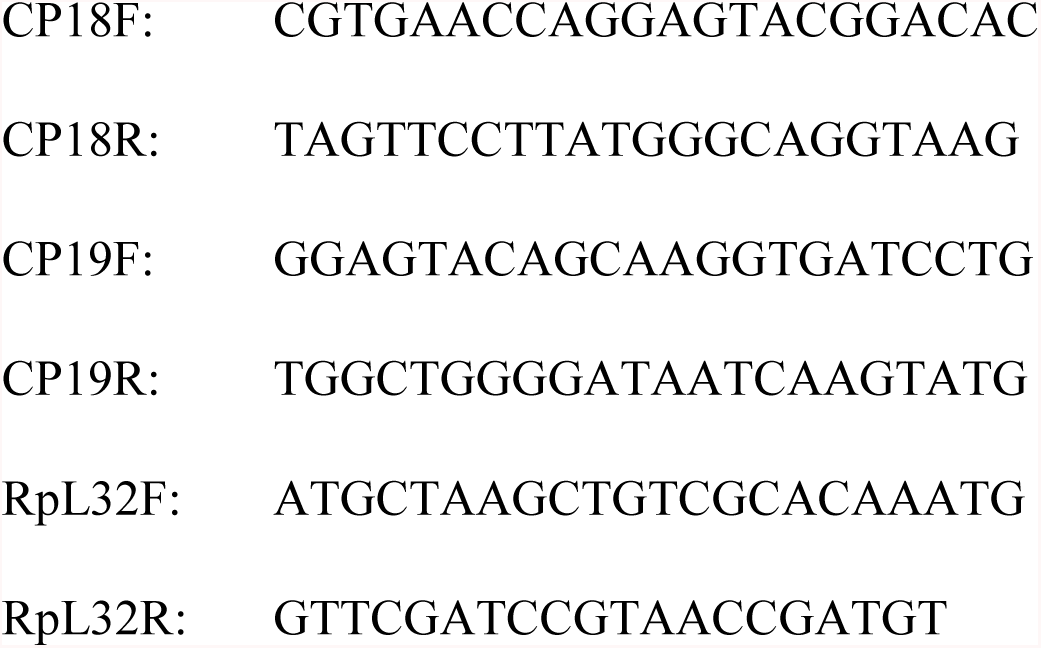

### Scanning electron microscopy (SEM)

Flies were placed on apple plates with wet yeast. The next day plates were changed to fresh ones with wet yeast and flies were let to lay eggs for 24h. Eggs were collected in TS-buffer (0.4% NaCl, 0.03% Triton X-100, Sigma), washed several times, and resuspended in PBS. Samples were fixed with 2.5% glutaraldehyde (Sigma) overnight and washed with NaPO4 buffer (pH 7.4). Dehydration was performed in ascending EtOH series (in 30% for 4h, in 50% EtOH for 4h, in 70% overnight, 96% for 2 × 4h, and 99.5% overnight) and critical point dried using Leica CPD300. Eggs were placed on SEM pins with carbon adhesive tabs (Electron Microscopy Sciences) and sputtered with platinum using Quorum Q150TS, turbomolecular-pumped high resolution coater with 30 mA sputtering current for 50s. Samples were imaged with FEI Quanta FEG Scanning Electron Microscope.

### Statistics

Statistical analyses were performed in Excel or OriginPro 2018. Shapiro-Wilk’s test was used to test the normality of the distributions of the measured values. For statistical comparisons, we used either Student’s t-test, with two-sample unequal variance or non-parametric Mann–Whitney test, with the significance level of 0.05.

### Data access

ChIP-seq and RNA-seq data are available under Gene Expression Omnibus accession number GSExxx from 1^st^ of August, 2018 onwards. Before this, please ask for a private token.

## Acknowledgements

We thank Paula Maanselkä for technical assistance and Dr. Joachim Urban for supplying UASp-Exp6 fly strain. Imaging was carried out at the Light Microscopy Unit (LMU) of the Institute of Biotechnology, SEM sample preparation as well as imaging at Electron Microscopy Unit of the Institute of Biotechnology, and next-generation sequencing at Biomedicum Functional Genomics Unit (FuGU) all of the University of Helsinki. This work was supported by the Academy of Finland, ERC, Sigrid Juselius and Jane & Aatos Erkko foundation grants to MKV.

## Author contributions

M.S. performed most of experiments and data analysis. H.M.M performed immunofluorescence experiments and quantification. B.P. performed qPCR and Western blotting, H.M.M., B.P. and M.H. egg counting and dorsal appendages length quantification. J.D., M.P. and V.H. designed and performed RanBP9 mutant fly generation. L.M. performed scanning electron microscopy. R.C.M. contributed to fly dissections. M.K.V. supervised experiments and data analysis. M.S, H.M.M. and M.K.V. wrote the manuscript. xM.S., H.M.M., M.P., V.H. and M.K.V. discussed the results and commented on the manuscript.

## Competing interests

None

